# Telomere shortening causes distinct cell division regimes during replicative senescence in *Saccharomyces cerevisiae*

**DOI:** 10.1101/2021.06.16.448683

**Authors:** Hugo Martin, Marie Doumic, Maria Teresa Teixeira, Zhou Xu

## Abstract

**Background:** Telomerase-negative cells have limited proliferation potential. In these cells, telomeres shorten until they reach a critical length and induce a permanently arrested state. This process called replicative senescence is associated with genomic instability and participates in tissue and organismal ageing. Experimental data using single-cell approaches in the budding yeast model organism show that telomerase-negative cells often experience abnormally long cell cycles, which can be followed by cell cycles of normal duration, before reaching the terminal senescent state. These series of non-terminal cell cycle arrests contribute to the heterogeneity of senescence and likely magnify its genomic instability. Due to their apparent stochastic nature, investigating the dynamics and the molecular origins of these arrests has been difficult. In particular, whether the non-terminal arrests series stem from a mechanism similar to the one that triggers terminal senescence is not known.

**Results:** Here, we provide a mathematical description of sequences of non-terminal arrests to understand how they appear. We take advantage of an experimental data set of cell cycle duration measurements performed in individual telomerase-negative yeast cells that keep track of the number of generations since telomerase inactivation. Using numerical simulations, we show that the occurrence of non-terminal arrests is a generation-dependent process that can be explained by the shortest telomere reaching a probabilistic threshold length. While the onset of senescence is also triggered by telomere shortening, we highlight differences in the laws that describe the number of consecutive arrests in non-terminal arrests compared to senescence arrests, suggesting distinct underlying mechanisms and cellular states.

**Conclusions:** Replicative senescence is a complex process that affects cell divisions earlier than anticipated, as exemplified by the frequent occurrence of non-terminal arrests early after telomerase inactivation. The present work unravels two kinetically and mechanistically distinct generation-dependent processes underlying non-terminal and terminal senescence arrests. We suggest that these two processes are responsible for two consequences of senescence at the population level, the increase of genome instability on the one hand, and the limitation of proliferation capacity on the other hand.

## Background

Human somatic cells have limited proliferation potential, a property that participates in ageing (1). The extremities of their chromosomes, the telomeres, preserve genome integrity but shorten with each cell cycle due to the end replication problem (2). Once they reach a critical length threshold, telomeres trigger a DNA damage response that permanently arrests cells in replicative senescence. Senescence is a long process, with human foetal cells dividing between 35 and 63 times *in vitro* before arresting, as measured in the first controlled analysis of the phenomenon (3). During senescence progression, cell division dynamics, arrest timing and death rates are very variable from cell to cell (4). Importantly, replicative senescence and its heterogeneity are associated with a specific increase in genomic instability, possibly contributing to cancer emergence with aging.

While the general mechanisms of telomere protection, telomere shortening and senescence onset have been studied for decades, the heterogeneity of senescence and its origins have received less attention. To address this aspect, we used the budding yeast model to explore the dynamics of cell divisions at the single-cell level (5, 6). *Saccharomyces cerevisiae* is a unicellular eukaryote that constitutively expresses telomerase, an enzyme that maintains telomere length homeostasis by preferentially elongating the short telomeres (7). Because we can experimentally repress telomerase expression, budding yeast is an excellent model to monitor the whole senescence process, from telomerase inactivation to senescence onset and cell death. Using this model, we and others showed that telomerase-negative cells often experience consecutive prolonged cell cycles or arrests from which they recover transiently before reaching terminal senescence (5, 8-13). These arrests are caused by the activation of the DNA damage checkpoint pathway, suggesting they stem from DNA damage. The succession of prolonged cell cycles relies on adaptation to DNA damage, a mechanism in which cells bypass checkpoint signalling to force mitosis (6). How these non-terminal arrests appear and contribute to the establishment of senescence, and whether they differ in nature from terminal arrests immediately preceding cell death, are unanswered questions. The way the cell handles the potentially resulting telomere instability likely contributes to the increase of genomic instability observed in telomerase-negative cells (6, 14, 15).

Because of the apparent stochastic nature of the early arrests, mathematical analysis and modelling are powerful approaches to describe them, test hypotheses on their nature and propose underlying laws governing their appearance. We previously built a telomere-shortening model that incorporated the asymmetric molecular mechanism of telomere replication (16, 17). Numerical simulations in cell lineages that do not experience non-terminal arrests, based on this model, suggested that a major source of variability of senescence onset is the wide length distribution of the shortest telomere in a population of cells.

Here, we exploit a more exhaustive set of experimental data derived from cell cycle duration measurements in individual yeast cell lineages captured in microfluidic chambers and tracked by time-lapse microscopy from telomerase inactivation to cell death (6). We mathematically describe the dynamics of appearance of non-terminal arrests. We then refine and apply a telomere-shortening model providing a mechanistic basis for their timing and heterogeneity. Finally, we argue that non-terminal arrests and terminal senescence correspond to two distinct regimes governed by different biological processes.

## Results

### Generation-dependent accumulation of cell cycle arrests is specific to the absence of telomerase

We took advantage of a data set we recently published (6), where the successive cell divisions of individual lineages of yeast cells were tracked using a microfluidics-based live imaging approach. The analysis of the resulting movies allowed the measurement of the duration of each consecutive cell cycle, with a 10 min time resolution. We previously highlighted the presence of abnormally long cell cycles in virtually all telomerase-negative cell lineages, which were interchangeably called cell cycle arrests because they relied on the DNA damage checkpoint (5, 6) (Fig. 1A and B). Long cell cycles were extremely rare in cells expressing telomerase and their frequency did not increase with the number of generations.

**Figure 1.**
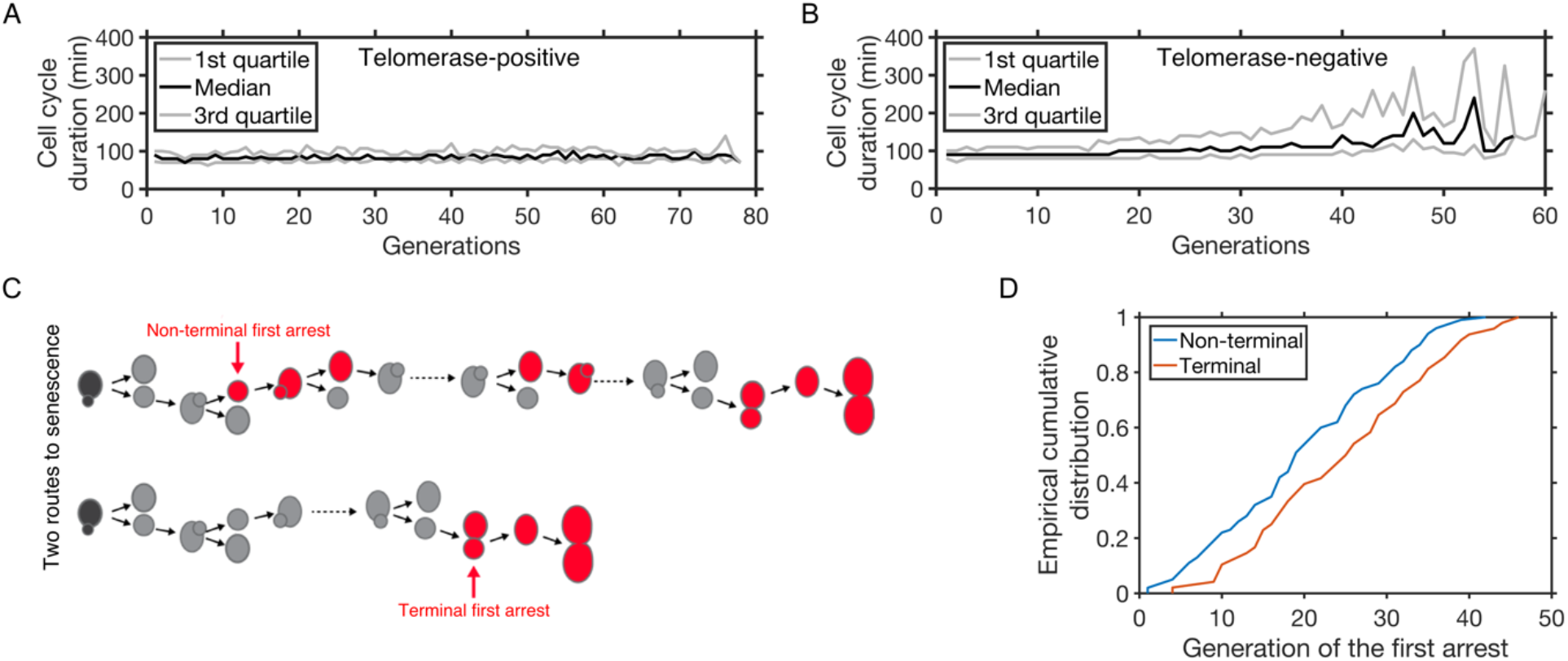
Telomerase inactivation leads to increasingly frequent cell cycle arrests. (A) Median cell cycle duration of cells from individual lineages as a function of generations in telomerase-positive cells. Gray lines represent the 1st and 3rd quartile. Generation count begins with image acquisition. (B) Same as (A) but in telomerase-negative lineages. Generation count starts with telomerase inactivation. (C) Scheme depicting two types of telomerase-negative lineages: (upper panel) the first arrest can be followed later on by at least one cell cycle of normal duration and is therefore “non-terminal”, or (lower panel) the first arrest can initiate the limited number of successive arrests characteristic of senescence and thus is “terminal”. (D) Cumulative distribution of the first appearance of terminal (red) and non-terminal (blue) arrests, with threshold *D* = 180 min.

We set a duration threshold *D* to define what was considered to be an abnormally long cell cycle. We observed that some of the abnormally long cell cycles, when appearing for the first time, were located at the end of the lineages and were never followed by a cell cycle of normal duration (Fig. 1C), consistent with first terminal senescence arrests being caused by the first telomere reaching a critical short length (16, 18, 19). Others, termed non-senescent, non-terminal or early arrests, were followed immediately or later on by at least one cell cycle of normal duration below *D* (Fig. 1C). In (5, 6), *D* was arbitrarily chosen to be the mean cell cycle duration of telomerase-positive cells plus three standard deviations. In the present work, unless otherwise stated, we also refer to long cell cycles or arrests using the same threshold *D*, which was computed from the telomerase-positive data set and was estimated to be 180 min. In addition, whenever possible, we challenged the robustness of the results with respect to this threshold. The fact that the first non-terminal arrests started generally earlier than the terminal arrests (Fig. 1D) suggested that they might be caused by a different mechanism and we therefore set out to characterize the law governing their appearance.

### The probability of appearance of the first non-terminal arrest increases over generations

We asked whether the first non-terminal arrest could be caused by a stochastic DNA damage event. Following this hypothesis, at each generation *j*, there would be a probability *p(j)* (0 ≤ *p(j)* ≤ 1) for an arrest to occur. Let *X* be the random variable describing the generation at which the first non-terminal arrest occurs. Then, for every positive integer k, we have:

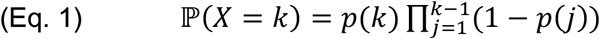

We first tested whether the event causing the arrest was independent of the number of generations, that is *p(j)* = *p* constant and *X* corresponds to Bernoulli trials. The generation of the first arrest would then follow a geometric distribution, of parameter *p* = 0.0465. However, a χ2 goodness-of-fit test rejected this hypothesis (p-value < 0.05), robustly with regards to the threshold *D* (Fig. 2A and Supplemental Fig. S1).

**Figure 2.**
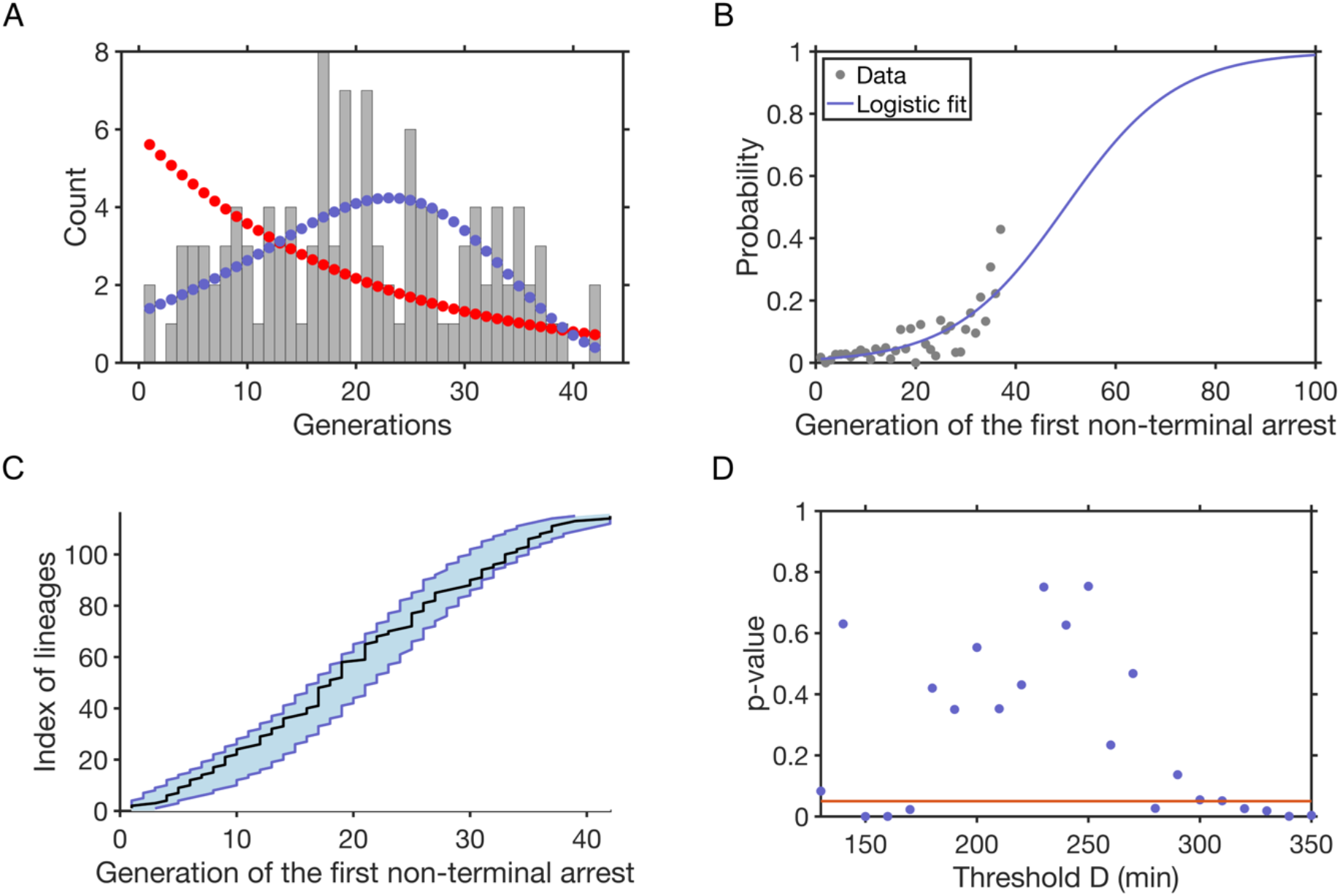
The stochasticity of appearance of the first non-terminal arrest is generation-dependent and best described by a variable Bernoulli parameter with an exponential growth. (A) Grey bars: distribution of the occurrence of the first non-terminal arrests with generations. Dots: Fit of the distribution with *p(j)* following the logistic function shown in (B) (blue dots) or being constant (red dots). (B) Grey dots: experimental frequency of occurrence of the first non-terminal arrest; blue line: logistic fit of *p(j)*, the probability of appearance of the first non-terminal arrest, calculated only for generations with more than 5 lineages left for computing. (C) Ordered generations of the first arrest from the experimental data (black line) or simulations (N = 1000) based on Bernoulli distribution of first arrest with an exponentially-growing parameter with the blue shaded area representing the 95% quantile. (D) p-values of χ2 goodness-of-fit tests of *X* corresponding to Bernoulli trials with a non-constant *p(j)* fitted in (A) as a function of the threshold *D*. The best parameters of (Eq. 2) are computed for each *D*. Red line represents p-value = 0.05. The hypothesis is not rejected for *D* between 180 and 270 min.

Plotting the fraction of lineages in which the first non-terminal arrest occurred at generation *j* out of all lineages that have not yet experienced an arrest at that point confirmed that a constant *p(j)* did not describe the experimental data accurately (Fig. 2B). Instead, *p(j)* increased in a non-linear, possibly exponential, manner. We tested this hypothesis and to ensure that 0 ≤ *p(j)* ≤ 1 for every *j*, we chose to fit the data with a logistic function that approximates an exponential growth for small values of *j* (Fig. 2B):

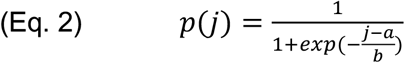

This approximation of *p(j)* allowed us to compute the generation of the first non-terminal arrest in simulated lineages. We found a good agreement between the simulation and the experimental data for the probability mass distribution function (Fig. 2A), in contrast to a geometric law, as well as for the cumulative distribution function (Fig. 2C), for a wide and biologically relevant range of *D* between 180 and 270 min (χ2 goodness-of-fit tests, Fig. 2D). Therefore, while the first non-terminal arrest seemed to occur stochastically, the probability of its appearance increased exponentially over generations, which might be consistent with a time-dependent or telomere-length-dependent damage as a plausible cause of the first arrest.

### A telomere-shortening-dependent model fits the timing of the first non-terminal arrest

We previously modelled the telomere shortening mechanism at the molecular level and found that a model with the shortest telomere in the cell reaching a signalling threshold explains the onset of the first terminal senescence arrests and their heterogeneity (16). Here, we asked whether a similar model of telomere shortening could also explain the first non-terminal arrest. Such a model would provide a mechanistic explanation for these arrests and would be consistent with the phenomenological description of the increase in their probability of occurrence over generations (Fig. 2B).

We therefore applied the telomere shortening model to look for a potential length threshold at which the first arrest would be induced. Each lineage started with an initial steady-state telomere length distribution simulated based on the length-dependent regulation of telomerase action. Telomeres were then allowed to shorten at each cell division, taking the asymmetry of telomere replication into account. We used the same approach as in (16, 17) and tested how the shortening dynamics of a linear combination of the lengths of the two shortest telomeres, that is *L*_*1*_ *+ α L*_*2*_ (where *L*_*1*_ and *L*_*2*_ represent the length of the shortest and second shortest telomeres, respectively, and *α* is a positive scalar) reaching a threshold *L*_*min*_ could fit the data. To do so, we simulated the senescence onsets of individual lineages according to (16, 17) and compared them to the experiments by minimizing the following error value, describing the distance between the simulation and the experimental data of the onset of the first arrest:

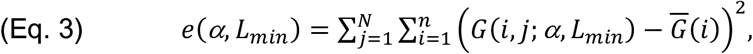

where *N* is the number of simulations, *n* is the number of experimental lineages (here *n* = 115) and for the *j*^*th*^ simulation, *G(i,j;a)* represents the *i*^*th*^ lineage out of *n* lineages, ordered from shortest to longest and simulated with the law of telomeric signalling *L*_*1*_ *+ α L*_*2*_ *= L*_*min*_, and 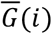 is the *i*^th^shortest lineage in the ordered set of the *n* experimental lineages.

Interestingly, as in (16) for the onset of senescence, the parameter *α* that minimized *e* was always 0, indicating that taking only the length of the shortest telomere into account, and not the second, led to the best fit. We therefore chose *α = 0* for the rest of the present work. The *L*_*min*_ that minimized *e* was 61 bp, larger than the value 19 bp found in (16), which was consistent with the fact that the first arrest appeared earlier than senescence onset (Fig. 1C). The simulated lineage data were ordered based on their length and the area representing the 95% quantile of *N =* 1000 simulations was represented (Supplemental Fig. S2). However, the simulations failed to capture the variation of the generation of the first non-terminal arrest (Supplemental Fig. S2). We concluded that a deterministic *L*_*min*_ was not a reasonable assumption given the variable nature of the first arrest.

At a molecular level, it would not be surprising that the signalling threshold for the shortest telomere might be probabilistic rather than deterministic (see discussion). To take into account this possibility, we chose *L*_*min*_ from a distribution defined by:

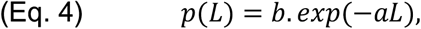

where *a > 0* and 0 < *b < 1* are parameters of the distribution. This distribution was chosen as a simple way to model a decreasing probability with only two parameters. At each division, one computes the shortest telomere length *L* and draws randomly, according to the probability *p(L)*, whether it signals the first arrest or not. (Eq. 4) shows that we assume that the probability of signalling senescence increases when *L* decreases. If a telomere happens to reach a length of 0 bp, which is a physiologically relevant situation (18), the arrest is immediately triggered regardless of this probability distribution.

Using numerical simulations (*N* = 1000), we fitted the parameters *a* and *b* on the experimental data and found the best fit to be *a* = 0.023 and *b* = 0.276, giving a signalling probability that is small if the shortest telomere is greater than 150 bp (*p*(150) = 0.009), and takes increasingly larger values as the shortest telomere decreases in length (Fig. 3A). This fit was in excellent agreement with the heterogeneity of the experimental data (Fig. 3B).

**Figure 3.**
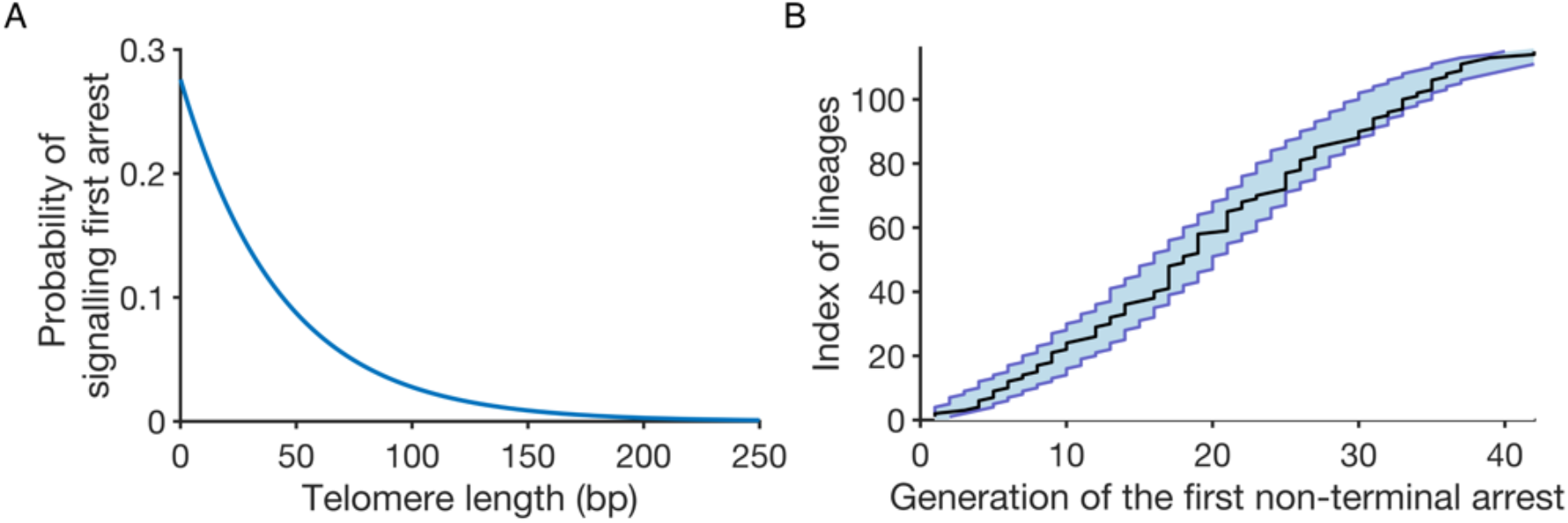
A telomere-shortening-dependent model with a probabilistic length threshold fits the timing of the first non-terminal arrest. (A) Probability density of the shortest telomere for signalling the first non-terminal arrest. (B) Ordered generations of the first non-terminal arrest from the experimental data (black line) or simulations (N = 1000) based on the probabilistic length threshold model with the blue shaded area representing the 95% quantile.

We conclude that a stochastic process or event depending on the length of the telomeres getting progressively shorter might explain the timing of occurrence of the first non-terminal arrest. Therefore, while the non-terminal arrests occurred slightly earlier than the terminal senescence arrests, both terminal and non-terminal arrests might be triggered by telomere-length-dependent processes (Fig. 1C and (16)). However, we do not exclude other sources of heterogeneity that might contribute to the observed timings. We speculate in the discussion about possible molecular mechanisms that would be consistent with the probability distribution we tested.

### The first non-terminal arrest is often followed by consecutive arrests

In the experimental data and our previous analyses, we observed the presence of consecutive long cell cycles corresponding to cells that managed to divide without repair, through adaptation (6). We also proposed that more generally, a first arrest increased the probability of occurrence of a second one (5). We thus closely examined the relationship between the first two non-terminal arrests. We first looked at the number of generations separating them and observed that the second arrest often immediately followed the first one (Fig. 4A). The overrepresentation of pairs of juxtaposed or closely positioned arrests (x = 1 or 2 in Fig. 4A) could not be explained by a random timing of occurrence of the second one as evidenced by the failure to fit a geometric distribution for a wide range of *D* (p-values of χ2 goodness-of-fit tests <0.05 most of the time; Supplemental Fig. S3).

**Figure 4.**
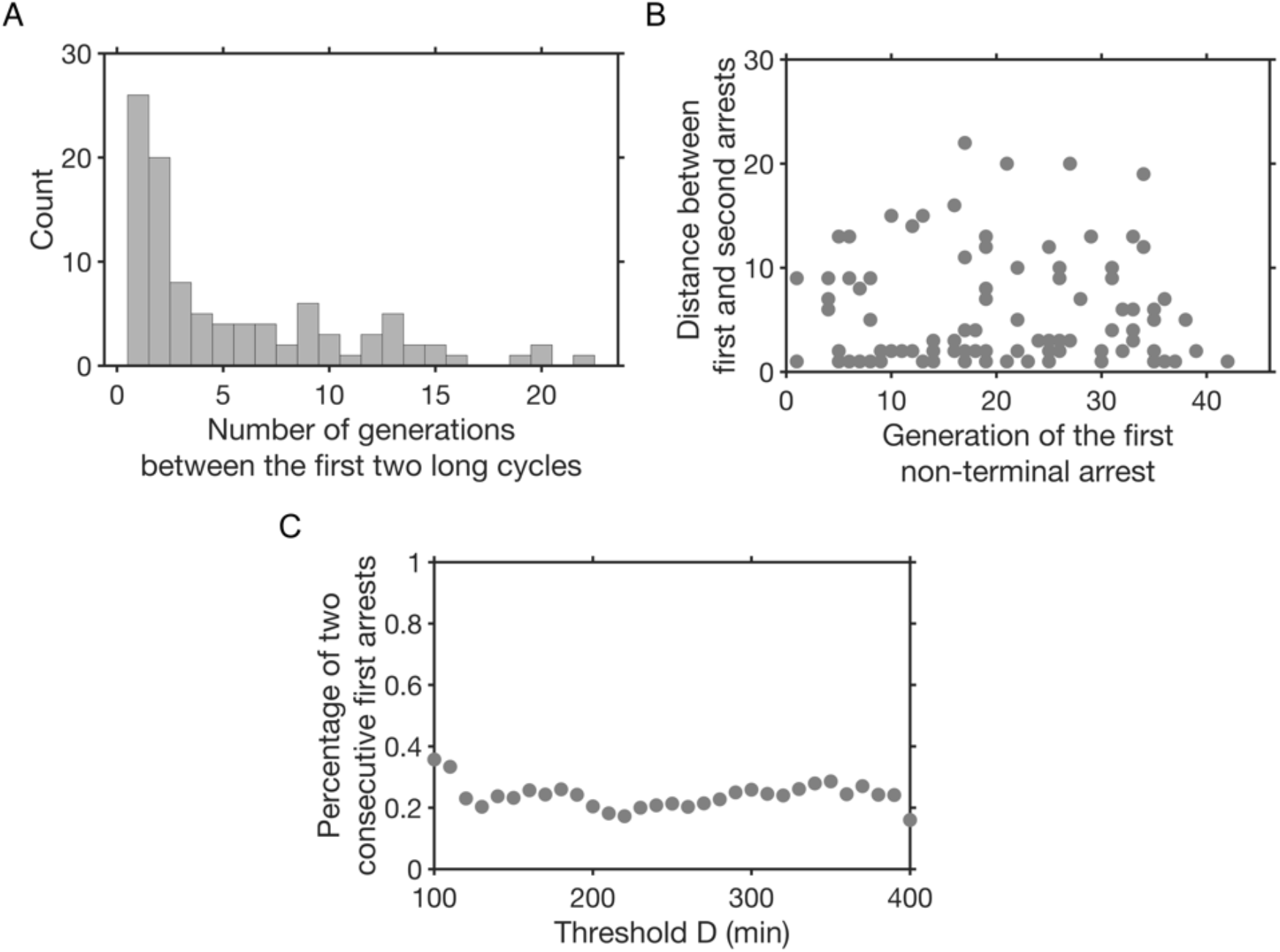
The first non-terminal arrests are often consecutive. (A) Distribution of the number of generations between the first two non-terminal arrests. (B) Number of generations separating the first two arrests as a function of the generation of the first arrest. (C) Fraction of lineages with consecutive first two arrests as a function of threshold *D*.

We then asked whether the generation of the first arrest affected the number of divisions before the next arrest. We plotted the number of divisions between the two first arrests against the generation of the first arrest and observed no significant correlation (Pearson correlation coefficient *R* = -0.029, *p*-value = 0.77, Fig. 4B). The overrepresentation of juxtaposed first two arrests were not just coincidentally due to setting the threshold *D* at 180 min since the fraction of lineages displaying this situation varied little as a function of *D* (Fig. 4C). These results are consistent with the cells dividing while transmitting an arrest-inducing defect, such as a signalling telomere.

### The non-terminal and senescent consecutive arrests correspond to two distinct regimes

Consecutive arrests are often associated with adaptation events and, after non-terminal arrests, the eventual return to cell cycles of normal duration suggests the repair of the initial damage (6). We therefore wondered if, for a sequence of non-terminal arrests, we could model the repair probability after a first arrest as constant, that is, we asked if the number of consecutive non-terminal arrests followed a geometric distribution (Fig. 5A). This hypothesis was not rejected (p-value > 0.05 for χ2 goodness-of-fit tests) for *D* between 140 and 330 min (Fig. 5A and B). For the standard *D* = 180 min, the estimated probability of repair was 0.65 per generation. We concluded that after the initial arrest, each further division is associated with a constant probability of repair and recovery.

**Figure 5.**
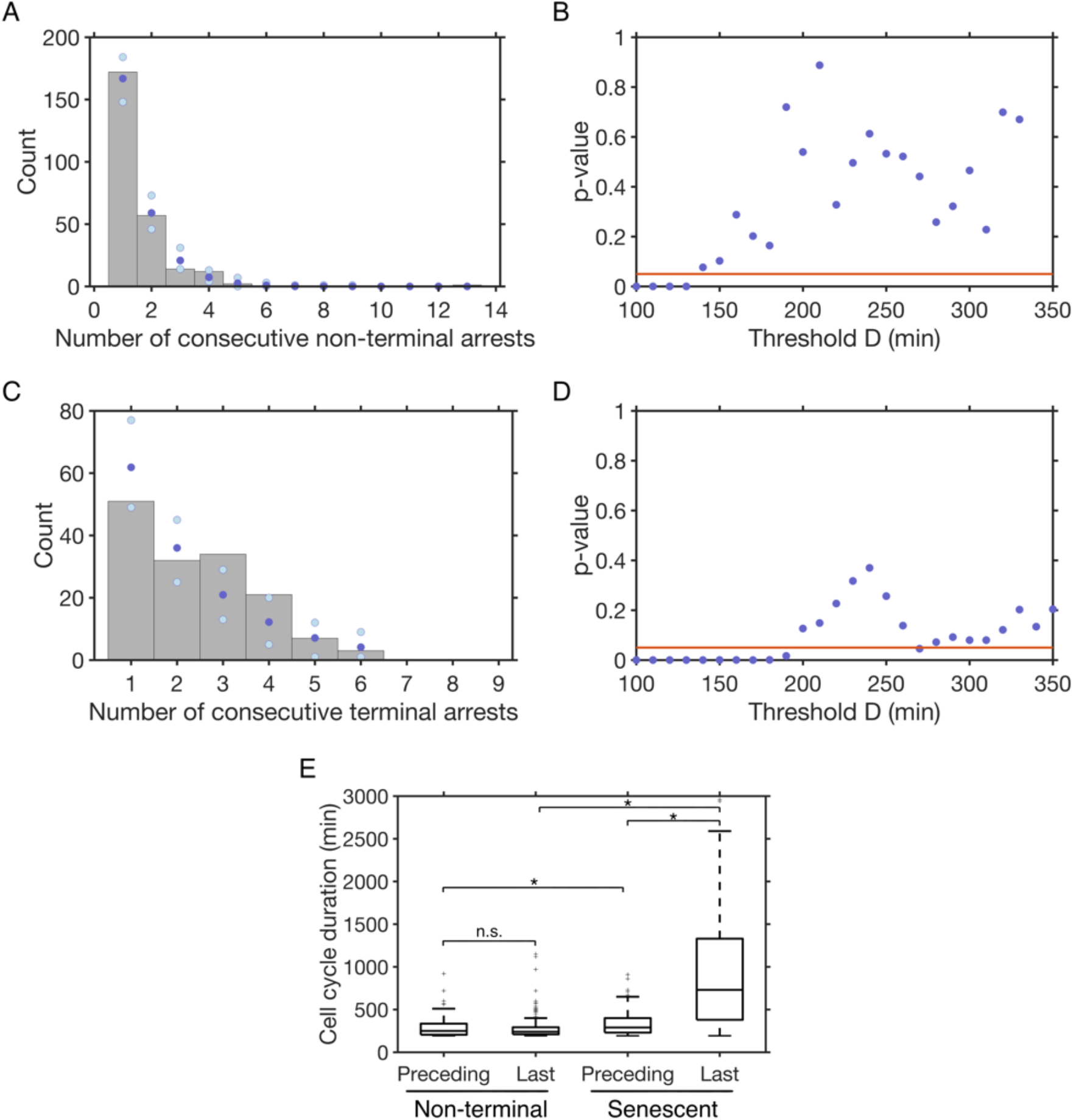
The distributions of the number of consecutive arrests reflect distinct regimes for non-terminal and terminal arrests. (A) Distribution of the number of consecutive non-terminal arrests for *D* = 180 min (grey bars) and fit with a geometric distribution (dark blue) with the 2.5% and 97.5% quantile shown in light blue. (B) p-values of χ2 goodness-of-fit tests for the number of consecutive non-terminal arrests following a geometric distribution, as a function of *D*. The hypothesis is not rejected for *D* between 140 and 330 min. (C-D) same as (A-B) for all terminal senescence arrests.

All lineages, whether they experienced non-terminal arrests or not, eventually enter senescence and undergo consecutive arrests before cell death. We asked whether each cell division in senescence would also have a constant probability of terminating the sequence of consecutive arrests. We found that this hypothesis of a geometric law was rejected for the standard *D* = 180 min (Fig. 5C and D). Therefore, the sequences of terminal and non-terminal arrests do not follow the same law, suggesting different underlying processes.

However, we observed that the hypothesis of geometric law for senescent arrests was not rejected for values of *D* between 200 and 350 min (Fig. 5D), values which were larger than for non-terminal arrests and more consistent with the timescale of persistent damage. Even at values of *D* where both non-terminal and terminal senescence arrests followed a geometric distribution such as 240 min, the parameter (between 0 and 1) defining the geometric laws and representing the probability of ending the consecutive arrests were different for the two processes: for example, 0.71 and 0.58 for the non-terminal and terminal senescence arrests, respectively, for *D* = 240 min. We then measured the difference in duration between non-terminal and senescence arrests and found that for a sequence of non-terminal consecutive arrests, the last long cell cycle had a duration on average similar to the preceding long cell cycles (medians of 250 min vs 240 min, respectively; Fig. 5E). In contrast, in senescence, the last cell cycle leading to cell death was much longer than the preceding ones (medians of 730 min vs 290 min, respectively; Fig. 5E). More generally, the cell cycle durations of non-terminal arrests were significantly shorter than for senescence arrests, even without taking the last arrest of the sequence into account (Fig. 5E).

Taken together, these results show that the dynamics of consecutive cell cycle arrests are kinetically different between non-terminal arrests and terminal senescence arrests.

The hypothesis is not rejected for values of *D* between 200 and 350 min. (E) Duration of the long cell cycles in a sequence of consecutive non-terminal or senescence arrests, separated into the last cell cycle in the sequence (“Last”) and the preceding ones (“Preceding”). * indicates *p*-value < 0.05 and “n.s.” non-significant for a Mann-Whitney U test.

## Discussion

In the present work, we mathematically describe and model the dynamics of appearance of cell cycle arrests in telomerase-negative yeast lineages. We distinguish the arrests arising for the first time near the end of a lineage, which are associated with the terminal senescence phase, from earlier non-terminal arrests, the origins of which are less well understood. Based on the analysis of the laws governing their dynamics, we argue that they correspond to two mechanistically distinct processes, both contributing to the complexity and heterogeneity of replicative senescence.

At first sight, the early arrests seemed to appear in a stochastic manner and could theoretically be explained by a global increase in replication stress or DNA damage, or by the disruption of non-canonical functions of telomerase, uncorrelated with the number of generations cells spent in the absence of telomerase. A strong argument that it was not the case came from the fact that similar experiments performed with cells with longer telomeres showed a considerable delay before these early arrests appeared (5). Here, the analysis of the probability of occurrence of the early arrests shows an exponential increase, confirming that they correspond to a generation-dependent process and rejecting all the aforementioned generation-independent hypotheses.

Another intriguing feature of these non-terminal arrests is that they are not isolated and often appear as a sequence of two or more consecutive arrests at a rate that exceeds by far the expectations even if we assume an exponential increase in the probability of appearance of the arrests. The geometric law governing the number of consecutive arrests for a range of thresholds *D* consistent with repair kinetics supports a model whereby early arrested cells undergo adaptation until the damage is repaired. Since adaptation promotes genome instability by forcing mitosis despite unrepaired DNA damage and lineages retain significant proliferation potential after non-terminal arrests, we speculate that these early arrests might contribute to the mutations and genome rearrangements observed in senescent cultures (8, 11, 14, 20). In contrast, cell cycles in the senescence phase end with an extremely long final arrest and cell death.

We propose that a signal dependent on the telomere-shortening mechanism is a plausible way to account for both the timing and heterogeneity of appearance of the first non-senescent arrests. Since the cells experiencing non-terminal arrests activate the DNA damage checkpoint, we speculate that the shortest telomere in these cells is in a signalling state that allows for repair or adaptation. In contrast, at a senescence arrest, the shortest telomere, while also triggering the DNA damage checkpoint, is likely unrepairable. One important parameter for the pathway choice between repair, adaptation and permanent arrest is the amount of exposed single-stranded DNA. It has been shown that the extent of resection and the recruitment of Rad51 and Rad52 increase as telomeres shorten (21-23). We can therefore imagine that different ranges of resection might lead to different cell fates and distinguish between non-terminal arrests and senescence arrests.

To model the appearance of non-terminal first arrests with generations after telomerase inactivation, we used a telomere-shortening model we previously built, in which the shortest telomere deterministically triggers the checkpoint arrest at a threshold length. The threshold that best fitted the experimental data is 61 bp, close to the 75 bp “recombination length” proposed in a recent work to promote activation of resection and initiate Rad51-dependent break-induced replication to elongate telomeres (24). However, this deterministic model does not capture the heterogeneity of the distribution of the appearance of the non-terminal arrests (Fig. S2), suggesting more complex events during the course of telomere shortening. We thus refined our model by allowing the length threshold for the shortest telomere to be probabilistic rather than fixed, which did not alter significantly the mean length threshold (62 bp) compared to the deterministic model. The reasoning behind this assumption is that telomere-length-dependent mechanisms, such as elongation by telomerase, DNA damage signalling or repair, are based on proteins bound to the telomere, including the telomeric Rap1 protein and its interacting partners. Therefore, a length measurement mechanism would only be as precise as the size of the binding site of proteins to DNA (25-28). For example, Rap1 is expected to bind on average every ∼19 bp on natural yeast telomeres. Because yeast telomere sequence is degenerated, the variability of the number of Rap1 molecules bound to different telomeres may be important. Besides, short telomeres bind the checkpoint sensor kinases Tel1 and then Mec1 when they become critically short, thus defining a two-step process to transition from non-signalling to pre-signalling and signalling states (18). We speculate that the recruitment of these kinases might also entail some level of stochasticity, which might depend on variable resection and exposure of single-stranded DNA. Another source of variability affecting the signalling capacity of telomeres could also come from the chromatin status of each telomere, as exemplified by the silencing and telomere position effect on transcription of subtelomeric genes and/or the histone levels near telomeres that are known to be affected in senescence (20, 29). Therefore, we suggest that the critical length threshold for the shortest telomere might be probabilistic. We show one implementation of a probabilistic threshold, which leads to a good fit of the experimental data. Nevertheless, we do not exclude alternative explanations involving other sources of heterogeneity such as various forms of DNA damage or stress associated with telomere shortening (e.g. replication stress, oxidative stress, mitochondrial dysfunction) that might alter the kinetics of cell divisions and arrests (30-34). We suspect that the *in vivo* process might be a combination of shortest telomere signalling-related and -unrelated heterogeneity.

## Conclusion

Overall, we propose that telomere shortening triggers a first non-senescent cell cycle arrest when the shortest telomere reaches a probabilistic length threshold. The cell then enters a sequence of arrests, which might rely on adaptation events until successful repair, with a constant probability with each cell cycle. The cell then resumes normal cell divisions until the shortest telomere again triggers a cell cycle arrest. After a succession of series of non-terminal arrests and normal cell divisions, the lineage finally enters senescence. In the senescence phase, by definition, no repair mechanism will allow cells to recover, except in the very rare cases of post-senescence survivors that are not modelled here. Instead, the cell can still adapt and undergo a limited number of divisions, until the last, and substantially longer, arrest before cell death. To conclude, given their kinetic and mechanistic properties, non-terminal and terminal arrests might contribute to different aspects of replicative senescence, with non-terminally arrested cells being able to propagate mutations induced by repair and adaptation, and terminally arrested cells being responsible for the loss of proliferation observed in senescent cell cultures.

## Methods

### Data curation

We use the data sets of telomerase-positive and telomerase-negative cell lineages from (6). We define senescence as the last series of consecutive cell cycles longer than threshold *D*, ending in cell death. When analyzing senescence arrests, lineages that are not dead at the end of the experiment are excluded, since we cannot determine whether they entered senescence or not. For telomerase-positive lineages, because the beginning and the end of image acquisition are not synchronized with cell divisions, we censor the first and last cell cycle of all lineages.

### Telomere shortening model

Telomere shortening is modelled following the mechanisms described in (17). Briefly, for each simulated lineage, the initial cell starts with 32 telomeres, the lengths of which are drawn from the steady-state distribution of telomere length as in (17, 19). At each cell division, in a given daughter cell, half of the telomeres shorten by a length corresponding to twice the shortening rate and the other half does not shorten, as posited by the mechanism underlying the end-replication problem. Only one of the two daughter cells is kept, thus defining a lineage. The telomere lengths in the lineages are then tested against different models for the signalling length threshold *L*_*min*_.

### Software and algorithm

All the minimization procedures and statistical tests were performed using Matlab 2020a and the CMAES algorithm (35) version 3.61 for Matlab.

## Declarations

### Ethics approval and consent to participate

Not applicable.

## Consent for publication

Not applicable.

## Availability of data and materials

The datasets and source codes supporting the conclusions of this article are available in GitHub: https://github.com/Hugo-Martin/telomeres

## Author contributions

M.D., M.T.T. and Z.X designed and supervised the study. H.M. and M.D wrote the mathematical model and performed the numerical simulations. M.D. and Z.X. wrote the first draft of the manuscript. All authors analyzed the data and revised the manuscript.

## Acknowledgements

We are grateful to Gilles Charvin for his critical reading of the manuscript.

