## Supplementary Information for "Telomere shortening causes distinct cell division regimes during replicative senescence in *Saccharomyces cerevisiae*"

**Short title:** Distinct cell division regimes leading to senescence

**Authors:** Hugo Martin[1], Marie Doumic[1*,corr], Maria Teresa Teixeira[2*] & Zhou Xu[3*,corr]

**Affiliations:**

[1]Sorbonne Université, JL Lions Laboratory, 75005 Paris, France

[2]Sorbonne Université, PSL, CNRS, UMR8226, Institut de Biologie Physico-Chimique, Laboratoire de Biologie Moléculaire et Cellulaire des Eucaryotes, F-75005 Paris, France

[3]Sorbonne Université, CNRS, UMR7238, Institut de Biologie Paris-Seine, Laboratory of Computational and Quantitative Biology, 75005 Paris, France

*: Co-last authors

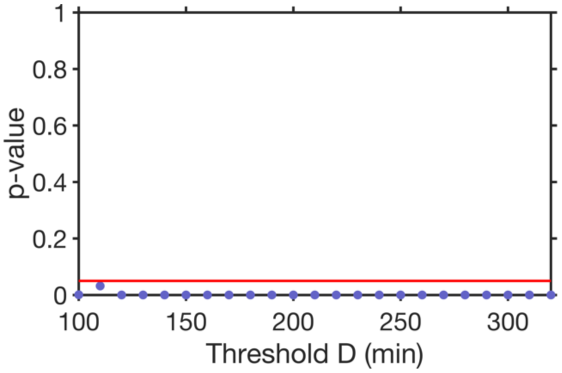


**Supplemental Figure S1.** The distribution of the timing of the first non-terminal arrest depends on the generation. Dots represent p-values of $\chi^2$ goodness-of-fit tests of the null hypothesis "*X* following a geometric distribution (with a constant parameter)" as a function of the threshold *D* (see fit in Fig. 2A, red dots). Red line represents p-value = 0.05. The hypothesis is rejected for all *D* tested.

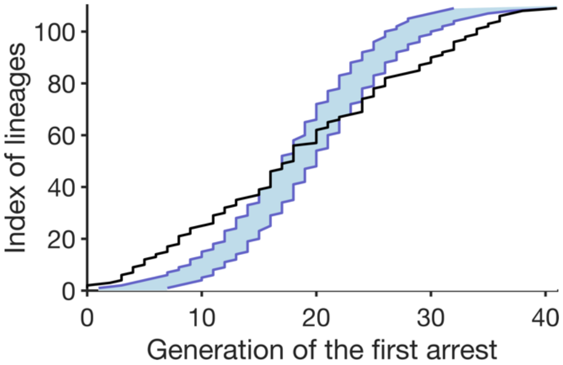


**Supplemental Figure S2.** A telomere-shortening model with a deterministic length threshold does not fit the experimental first non-terminal arrest data. Ordered generations of the first non-terminal arrest from the experimental data (black line) or simulations based on the deterministic length threshold model (N = 1000), with the blue shaded area representing the 95% quantile.

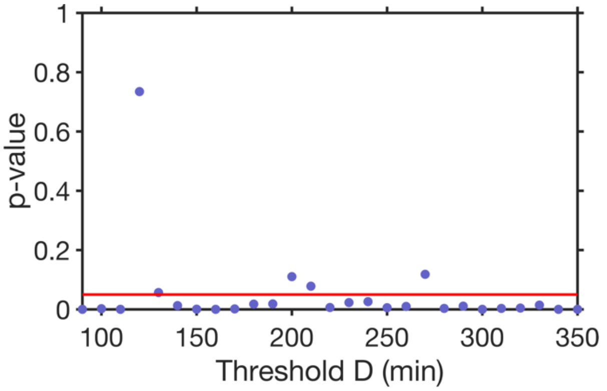


**Supplemental Figure S3.** The overrepresentation of pairs of juxtaposed or closely positioned arrests is not compatible with a random distribution of arrests. Dots represent p-values of $\chi^2$ goodness-of-fit tests for the distribution shown in Fig. 4A being a geometric distribution as a function of the threshold *D*. Red line represents p-value = 0.05. The hypothesis is rejected for most *D* tested.
